# Accuracy and precision of citizen scientist animal counts from drone imagery

**DOI:** 10.1101/2020.12.03.409649

**Authors:** Sarah A. Wood, Patrick W. Robinson, Daniel P. Costa, Roxanne S. Beltran

**Affiliations:** Department of Ecology and Evolutionary Biology, University of California Santa Cruz, 130 McAllister Way, Santa Cruz, California, 95060, USA; Institute of Marine Sciences, University of California Santa Cruz, 115 McAllister Way, Santa Cruz, California, 95060, USA

**Keywords:** Community science, seals, sea lions, drones, UAS, marine mammal

## Abstract

Repeated counts of animal abundance can reveal changes in local ecosystem health and inform conservation strategies. Unmanned aircraft systems (UAS) such as drones are commonly used to photograph animals in remote locations; however, counting animals in images is a laborious task. Crowd-sourcing can reduce the time required to conduct these censuses considerably, but must first be validated against expert counts to measure sources of error. Our objectives were to assess the accuracy and precision of citizen science counts and make recommendations for future citizen science projects. We uploaded drone imagery from Año Nuevo Island (California, USA) to a curated Zooniverse website that instructed citizen scientists to count seals and sea lions. Across 212 days, over 1,500 volunteers counted animals in 90,000 photographs. We quantified the error associated with several descriptive statistics to extract a single citizen science count per photograph from the 15 repeat counts and then compared the resulting citizen science counts to expert counts. Although proportional error was relatively low (9% for sea lions and 5% for seals during the breeding seasons) and improved with repeat sampling, the 12+ volunteers required to reduce error was prohibitively slow, taking on average 6 weeks to estimate animals from a single drone flight covering 25 acres, despite strong public outreach efforts. The single best algorithm was ‘Median without the lowest two values’, demonstrating that citizen scientists tended to under-estimate the number of animals present. Citizen scientists accurately counted adult seals, but accuracy was lower when sea lions were present during the summer and could be confused for seals. We underscore the importance of validation efforts and careful project design for researchers hoping to combine citizen science with drone imagery.

## 1. Introduction

Abundance is a critical metric for wildlife conservation, management, and policy [1-3]. This is especially true for seals and sea lions (hereafter, pinnipeds) that spend much of the year foraging at sea and can only be counted when they haul out on land to rest or reproduce [4, 5]. Yet counts from traditional methods such as visual ground surveys consistently underestimate true animal abundance [6-8], especially for dense groups of animals in rugged, inaccessible terrain [9]. Ground counts also disturb resting animals which can exacerbate underestimation. Pinnipeds have also been counted in photographs taken from manned aircraft [8, 10, 11], but these counts are costly and tend to be cost-prohibitive for longitudinal studies.

Unmanned aircraft systems (UAS; hereafter, drones) can provide high-resolution photographs and geospatial data for wildlife surveys [12], individual identification [13], and photogrammetry [14]. Despite being limited by weather and the potential to disturb animals if altitude recommendations are not followed [15], drones have been utilized to count various species [12]. Benefits of drone censuses include: more animals are counted, remote locations can be accessed, they are relatively inexpensive, they require less personnel in the field, minimize animal disturbance, and they create a permanent photographic archive [7, 16, 17]. However, counting animals manually from drone photographs can be a tedious and time-consuming task. While these technological advancements have opened new doors for the future of population ecology [12, 16–18], methods for quickly processing the resulting imagery have lagged.

Citizen science is a potentially useful tool that involves recruiting volunteers from the public without prior experience to complete clearly outlined tasks [19]. Despite concerns of bias and volunteer skill, citizen science has been used to gather and process large datasets for over a decade [20, 21], including for research on abundance and distribution of marine mammals [22–25]. Crowd-sourced science requires an up-front investment of time and labor to create a training system for new volunteers [26, 27]. However, once trained, citizen scientists can process data at no cost, leaving project managers to focus their time and funds on other aspects of the project [27]. Long-term monitoring projects from drone surveys or satellite imagery could greatly benefit from citizen science’s data processing capabilities [28].

The pinniped colonies at Año Nuevo Island offer an ideal opportunity to validate wildlife drone censuses. Pinnipeds are present throughout the year, and the island is easily accessible via drone and inflatable boat from the mainland and lacks vegetation, so animals are easily spotted. For our validation study, we focus on pinnipeds, specifically northern elephant seals (*Mirounga angustirostris),* harbor seals (*Phoca vitulina*), Steller sea lions (*Eumetopias jubatus),* and California sea lions (*Zalophus californianus)*, because their well-documented haul out patterns provide a useful comparison to past counting methods. We used the Zooniverse platform (zooniverse.org) to create a citizen science project titled Año Nuevo Island Animal Count, where volunteers were trained and instructed to count pinnipeds in drone photographs. Our objective was to explore the benefits and drawbacks of using citizen science to census pinnipeds in drone images. Here, we detail our methods for creating a citizen science project and use various analyses to evaluate citizen science accuracy. We also document our strategies for volunteer engagement, specific tutorials, and supplemental materials recommended by Swanson, Kosmala (29). If citizen scientists can accurately complete a census project, drones could substantially reduce the time and labor required for population surveys.

## 2. Materials and Methods

### 2.1. Study Site and Drone Flights

We conducted drone flights above Año Nuevo Island (37.1083°N, 122.3378°W), located on the West Coast of California, USA, using recommended best practices [30]. Año Nuevo Island is a rookery for many marine bird species and a breeding site for multiple pinniped species. The cliffs, flat terraces, and surrounding kelp forests provide diverse habitat and food supply for thousands of animals to rest, feed, and reproduce. Due to extensive restoration efforts on the island, bird nests are mostly confined to upper terraces, and pinnipeds occupy the edges, although species in the sea lion family *Otariidae* (*Z. californianus and E. jubatus*) often climb onto the terraces. From January to March, elephant seals haul out on the island’s beaches for the annual breeding season. The seals will then leave on a foraging migration, returning from April until August, dependent on their age and sex, to undergo their catastrophic molt to shed their fur and skin [31, 32]. California and Steller sea lions give birth beginning in June and are abundant on the island year-round because they undertake shorter foraging trips than elephant seals [33, 34].

Two consumer-level drones were used for the project: the Phantom 3 Advanced and the Mavic 2 Zoom (SZ DJI Technology Co., Ltd., Shenzhen, China; cost ~$1,000 USD each). Drone flights were conducted roughly every two weeks from July 2017 to July 2019 (N=60 flights, Table S1) depending on rain, wind, and swell conditions that would compromise drone flight safety or photo quality. To minimize animal disturbance, we launched the drone from the mainland and flew over the 1 km ocean channel to the island at 40 meters above sea level. We undertook drone flights early in the morning (typically 7:00-8:00 am) to maximize the number of animals present on the beaches due to cooler weather. The Litchi application (VC Technology Ltd. London, England) was used to produce a standardized flight path over the island (Figure 1A). Photographs were collected approximately every 2 s along the flight path at a speed of 20 kph. To improve each image’s location accuracy, we used a hand-held GPS unit (accuracy set to 5 meters) to measure the latitude and longitude of five locations on the island that were used as 2-dimensional ground control points (hereafter, GCPs).

**Figure 1.**
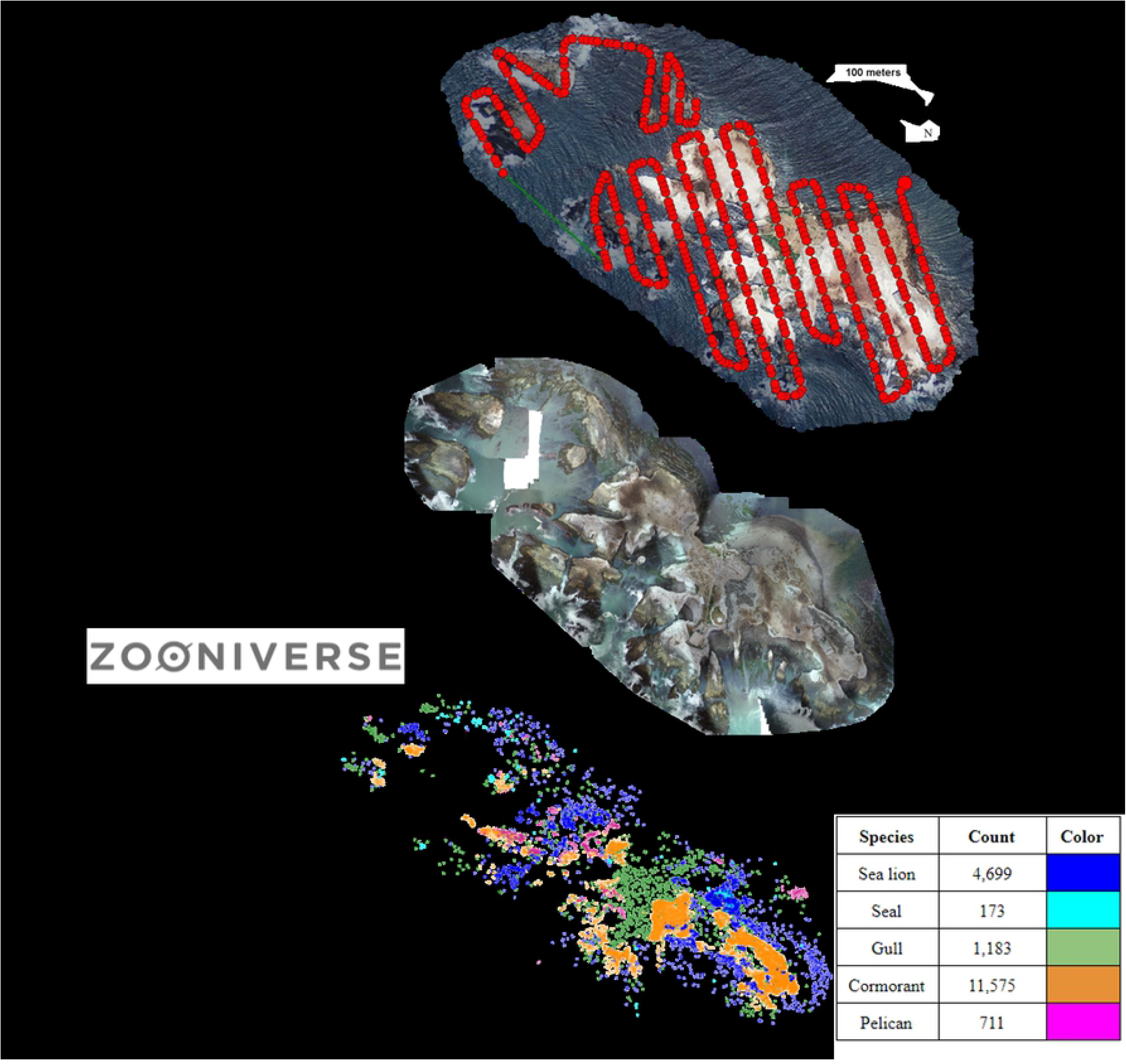
The complete workflow to compile individual images from a July 5, 2019 drone flight into an orthomosaic for citizen science and expert counts (18,366 animals).

### 2.2. Photograph Processing

The ~500 drone photographs per flight were uploaded into the photograph stitching software Pix4d (Pix4D S.A., Prilly, Switzerland) to produce a whole-island mosaic (Figure 1B). We used the 3D maps template with the standard system’s settings but changed the coordinate system to WGS 84 and manually entered the five GCPs. After processing, we rejected mosaics with poor score reports indicating insufficient photograph matching (N=8 flights). We then used *R* version 3.6.1 (R Core Team, 2019) to convert the complete island mosaic from a .tif to a .jpeg file using the *R* package *jpeg* with quality set to one. Next, we cut the .jpeg into a standardized 700 x 700-pixel jpegs using the *R* package *drones* (~700-1,000 tiled photographs per flight, hereafter “tiles”). We removed any photographs smaller than 13 kb because these only contained white space.

### 2.3. Citizen Science Website Development

We uploaded image tiles into a custom project on the free citizen science platform Zooniverse (Figure 2) and curated content about the team members, species natural history, preliminary results, frequently asked questions, and a discussion board. Each day, we answered questions and posted announcements on the discussion boards to keep volunteers engaged. Website design and volunteer engagement was a critical part of our strategy (see Supplemental Material). For the animal counting task, we used the “Drawing” program, which records the category (e.g., a pinniped species) and x and y coordinates of each mark added by volunteers. Prior to counting, volunteers were given a short tutorial with example photographs. Double-counting of animals in adjacent photographs was avoided by instructing volunteers to mark each animal only if its head was visible in the photograph.

**Figure 2.**
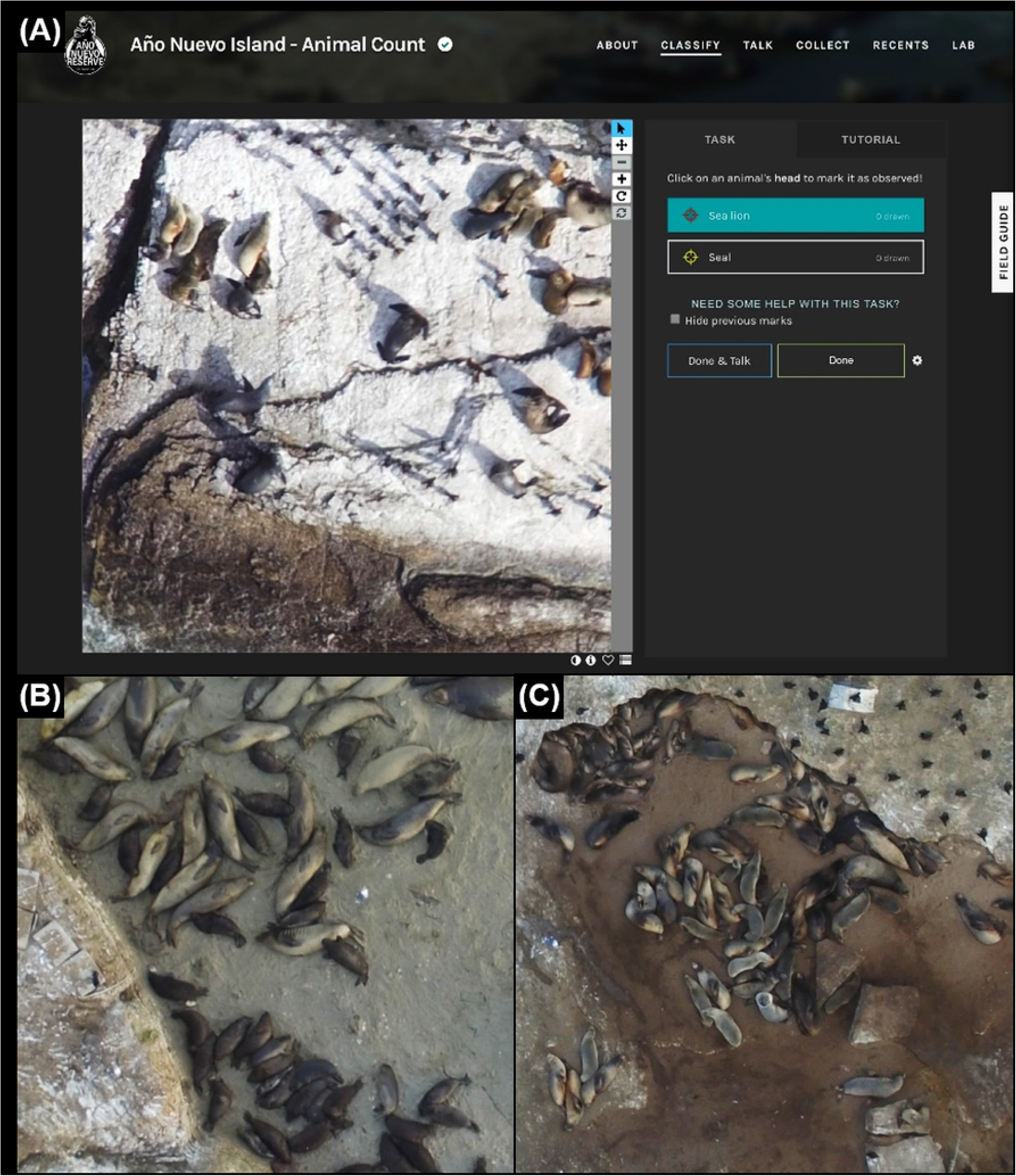
**(A)** A screenshot from the “classify” tab of the citizen science website (sealcount.com) with additional drone photographs of **(B)** elephant seals and **(C)** sea lions for illustrative purposes. Users are asked to select “seal” or “sea lion” and click once on each animal’s head to count it. All volunteers have access to a tutorial and field guide with detailed instructions for animal identification.

### 2.4. Beta Test Accuracy

Zooniverse requires a beta test with the full instructional tutorial and a small subset of photographs before publishing a project. We selected and uploaded 100 tiles that included a mix of pinnipeds and birds (to assess the most frequently misidentified species) and a small subset containing no animals (to assess which objects such as rocks were most frequently misidentified as animals). Beta testers were asked to count animals and subsequently provide feedback and suggestions for project improvement. Each photograph was considered complete after ten beta tester volunteers counted the seals, sea lions, and birds. We inspected beta tester counts for accuracy of counts and the number of counters needed to maximize consistency. The beta tester volunteers provided valuable feedback regarding the difficulty of identifying multiple species and the tutorial’s complexity. Many mistakes (e.g., counting only pinnipeds but no birds) were consistently repeated.

Based on our visual inspections and feedback results, we removed birds from the workflow and increased our requirement from 10 to 15 volunteers per photograph to allow us to determine which of several algorithms would yield the most accurate estimates. To expedite the data collection process, we eliminated photographs marked empty (i.e., no animals) by seven volunteers. Due to the difficulty in identifying pinniped species, volunteers on our citizen science website were asked only to classify animals based on pinniped family (seals in the family *Phocidae* included *P. vitulina* and *M. angustirostris* and sea lions in the family *Otariidae* included *Z. californianus* and *E. jubatus*).

### 2.5. Full Count Validation

We uploaded 4,074 photograph tiles from five drone flights that spanned winter and summer for the full validation effort. We launched the project at www.sealcount.com, actively engaged volunteers, and advertised the site through social media and news agencies, local educational organizations, and middle and high school classrooms. We also hosted two week-long counting contests during the first and last months of the project, where volunteers could win stickers or stuffed animal prizes for counting the most images over the week. After 212 days of citizen scientist counting, we downloaded and analyzed the count data. To collapse the multiple citizen scientist counts per tile into a single count, we used six algorithms: Mean without the two lowest values (Mean[3:Max]), Mean, Mean without the two highest values (Mean[1:Max-2]), Median without the two lowest values (Median [3:Max], Median, and Median without the two highest values (Median [1:Max-2]). Seal and sea lion count data were evaluated separately.

We also conducted expert counts on images of the same resolution as those presented to the volunteers. To create expert counts, each animal was classified using the cell count function in ImageJ as: sea lions, seals, gulls, cormorants, or pelicans. Only northern elephant seals were counted during the two winter flights because they made up nearly all pinnipeds on the island. After using the algorithms to create a single citizen science count per tile, we compared the citizen science counts and expert counts of elephant seals and sea lions by calculating relative error as the absolute value of the difference between the citizen science count and the expert count, divided by the expert count. The percent error calculations were repeated in two ways: (1) for each photo and (2) for each unique number of citizen scientists that counted each photograph (Table S2). Because percent error calculations are not possible for photographs in which the expert counted zero animals (i.e., the denominator is zero), we also calculated the proportion of photographs that were classified as true negatives (both citizen scientists and the expert counted no animals), true positives (both citizen scientists and the expert counted at least one animal), false negatives (citizen scientists counted no animals but the expert counted at least one animal) and false positives (citizen scientists counted at least one animal but the expert counted no animals) (Figure S2). A paired Wilcoxon Rank Sum test was used to quantify differences between citizen scientist counts and expert counts because count data were not normally distributed. Finally, we created expert counts for all elephant seals in 52 drone flights from July 2017-July 2019 to assess the elephant seal’s overall abundance on the island. Citizen science counts were overlaid into the 2017-2019 expert counts to assess trends in abundance visually. Data are available at https://doi.org/10.7291/D1J66X.

## 3. Results

### 3.1. Citizen Scientist Website Response

Between the website launch and the download of data for this analysis (August 7, 2019, and March 7, 2020, respectively) more than 1,500 volunteers counted ~94,000 tiles. On average, 2,500 photographs were counted each week. The most frequent counts occurred during our launch week (12,000 photographs counted) and counting contests (7,313 photographs counted during the sticker contest and 10,981 photographs counted during the stuffed animal contest). The contests were critical for reaching our count goals. Considerable upticks in counts also occurred during public outreach events and classroom visits, especially when guests could count during the events on individual computers. Approximately 30,000 images were counted by volunteers without a Zooniverse account and 60,000 counted by those with an account. Throughout the project, the average volunteer counted 59 photographs in total, with a handful of volunteers counting over 1,000 photographs and a maximum of 5,153 images counted by one individual (Figure S1). The top three volunteers completed 24% of the images counted by volunteers logged in to their Zooniverse accounts, or 15% of the total photographs counted by all volunteers.

### 3.2. Full Count Validation

For both seals and sea lions, percent error was lower for Median algorithms as compared to Mean algorithms (Figure 3). The single best algorithm was Median without the lowest two values (Median[3:Max]; 43% error for seals and 30% error for sea lions), suggesting that citizen scientists tended to under-estimate the number of sea lions present. This relatively large percent error was likely because not all photographs reached their retirement limit of 15 counters, and some photographs were counted by very few counters (Figure 3). Percent error decreased as the number of classifiers increased (Figure 3). The relationship between the number of counters and percent error was consistent across seals and sea lions, reaching an asymptote of percent error at around 11 citizen scientists; however, the relationship was more variable for seals (Figure 3). Because sea lions were far more common in photographs than seals (Figure 1), we hypothesize that the high error and variability in seal counts resulted from misclassification of seals as sea lions. Additionally, the small number of individuals in each photograph (Median=2, Mean=6, SD=12 for seals; Median=5, Mean=12, SD=19 for sea lions) meant miscounting one or two individuals led to a large error.

**Figure 3.**
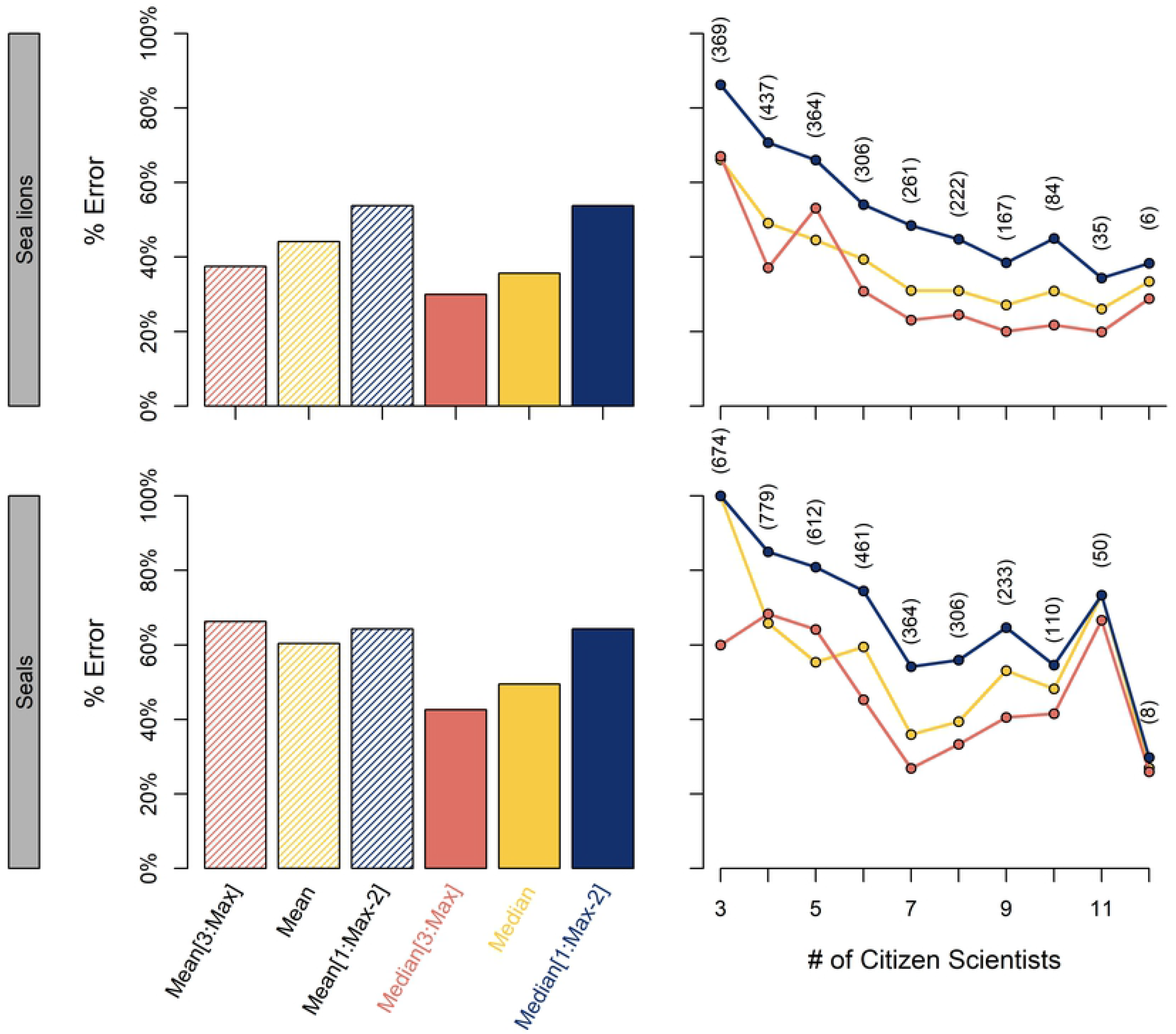
Comparison of percent error for (left) six algorithms for all photographs and (right) based on the number of citizen scientists that counted each photograph (number of photographs provided in parentheses). Algorithms using the Median consistently outperformed those using the Mean, especially for elephant seals. Because scientists tended to underestimate sea lion abundance, the most accurate counts were obtained using many repeated counts and removing the lowest two values before calculating the Median.

Across all species and median algorithms, approximately 80% of photographs were classified correctly in terms of the presence or absence of animals (Figure S2). For sea lions, 42-46% of photographs contained true negatives, 34-40% of photographs contained true positives, 9-12% of photographs contained false positives, and 6-12% of photographs contained false negatives. For seals, 76-81% of photographs contained true negatives, 3-5% of photographs contained true positives, 2-5% of photographs contained false negatives, and 12-17% of photographs contained false positives. In other words, if citizen scientists were to make a mistake in classifying seals, they were more likely to mark a non-existent animal than to miss an existent animal. The low proportion of true positives for seals is due to their relatively limited abundance and small spatial range on Año Nuevo Island.

The relationship between counts from citizen scientists and the expert for each photograph was relatively good (Figure 4). More variance could be explained in the sea lion counts (*R^2^* range from 0.82 to 0.93) compared to the seal counts (*R^2^* range from 0.64 to 0.81). The slopes of all algorithms for elephant seals were less than one, suggesting that underestimation was more common when many animals were present in photographs (Figure 4).

**Figure 4.**
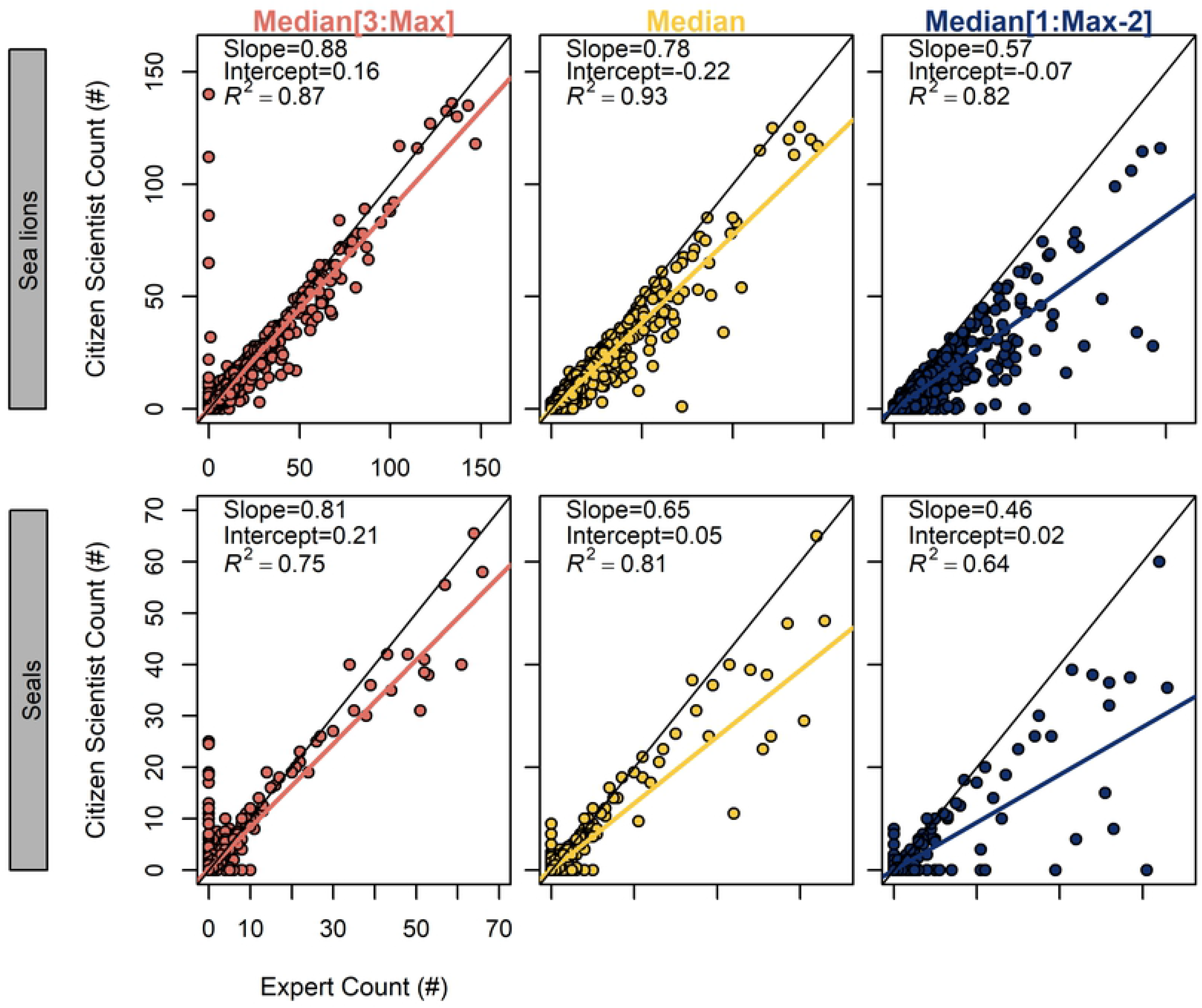
Comparison of raw citizen scientist counts against expert counts for three algorithms. Citizen scientists tended to underestimate the actual number of pinnipeds present on Año Nuevo Island, as evidenced by the slopes <1 for all algorithms and species except for Median[3:Max] with elephant seals (bottom left). Colored lines represent a linear model for each algorithm and species combination, whereas black lines represent 1:1 relationship between citizen scientist count and expert count.

After summing citizen scientist counts to come up with a single count for each algorithm and drone flight to compare with the expert count, we discovered that citizen science counts had variable accuracy after all tiles were combined (Figure 5). Specifically, the percent error of citizen science counts compared to expert counts was 9% for sea lions and 46% for seals across all dates (Figure 5). The seal counts were far more accurate during the winter flights (6% error for both) than the summer flights (54%, 69%, and 95% error), likely due to the presence of sea lions that were misidentified as seals. The higher percent error for seals compared to seals was likely due to their overall lower abundance (Figure 5).

**Figure 5.**
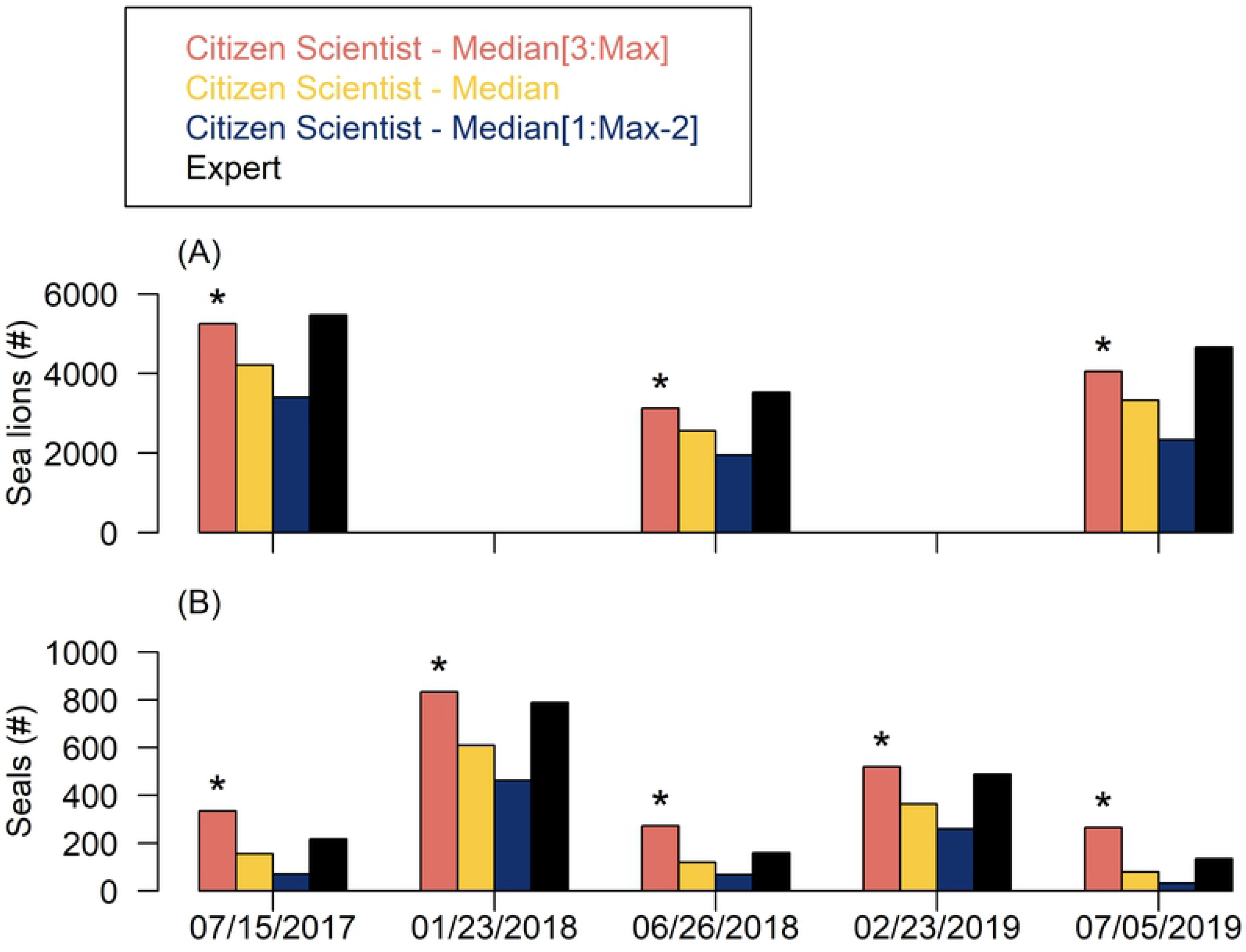
Counts for each drone flight date by an expert as compared to the three Median algorithms for summarizing citizen science counts. Sea lions were not counted during the two winter flights due to extremely low abundance. An asterisk denotes the citizen science algorithm (Median) that best matches the expert counts across all flights.

Because citizen science counts were comparatively slower and could only be obtained for 5 out of 52 flights, detailed elephant seal abundance trends could not be observed, unlike the high-resolution expert counts (Figure 6). The bi-weekly expert counts of elephant seals from 2017-2019 showed the expected fluctuation in abundance across various life history events, with the least number of animals present during foraging seasons (summer and late winter), and the most animals present during the breeding haul-out (winter) and molting haul-out (late spring) (Figure 6). Peak abundance occurred during the molting haul-out.

**Figure 6.**
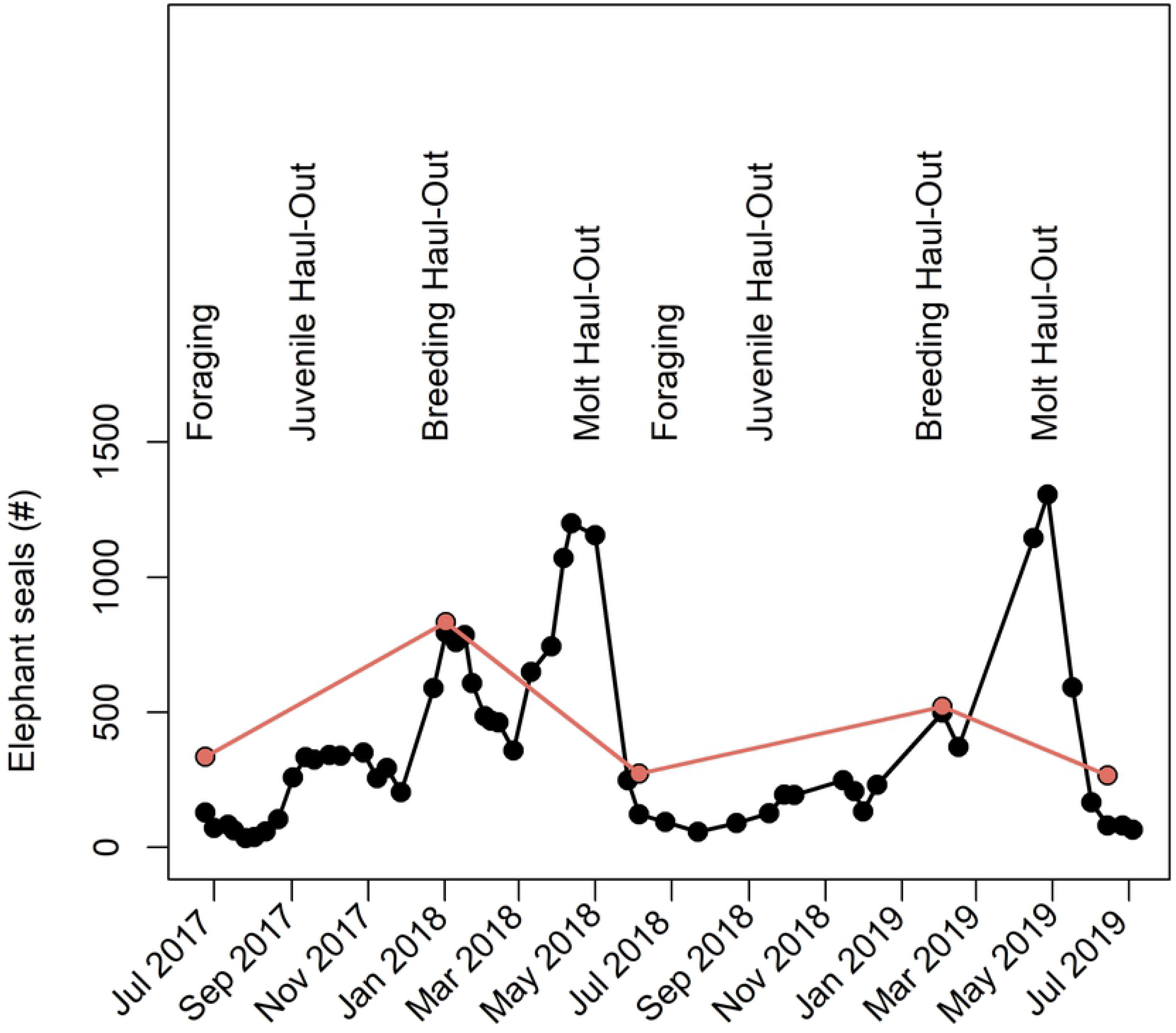
The total number of elephant seals counted by experts (black points) as compared to citizen science counts using the Median[3:Max] algorithm (pink points).

## 4. Discussion

We found that citizen scientists could accurately identify sea lions in most circumstances but had more difficulty identifying seals, especially when sea lions were present. Generally, citizen scientists tended to underestimate the abundance of animals on Año Nuevo Island. One possible explanation for this underestimation is that the small size and dark coloration of newborn seal and sea lion pups could cause citizen scientists to miss them, mistake them for rocks, or mistake them for a different species. Proper precautions must be taken to use citizen science as an alternative to ground surveys and expert counts. We recommend simplifying as much as possible to single demographic categories (e.g., only seals, only sea lions, or only pups). A critical consideration is that ground surveys are nearly impossible for many locations with rugged terrain, including Año Nuevo Island. The sheer number of animals makes an accurate count nearly impossible (i.e., scientists estimate sea lions by visually summing groups of ~500 animals).

As highlighted in other studies [29, 35], we found that repeat counts (11+ counters per image) were necessary to improve accuracy (Figure 3). More citizen scientist counts resulted in lower error; however, this requirement for repeated counts coupled with a moderate participation rate made citizen scientist counts prohibitively slow. Project managers should interact with volunteers as much as possible, as more photographs were counted when we held events or contests, and a large proportion of total classifications were done by a small group of citizen scientists (Figure S1). Finally, creating a thorough and easy-to-read tutorial is critical to project success. Tutorials must be concise and only give information pertinent to the task volunteers will perform. Including additional information elsewhere for volunteers to explore is recommended, but the tutorial is the crucial point at which volunteers decide to participate [36].

When beginning our project, we underestimated the importance of ensuring that the project was not too complicated. Using only expert counts for a census would yield more accurate results overall but would require a significant amount of labor, especially for large datasets. In comparison, citizen science allows large amounts of data to be collected inexpensively and in a relatively short amount of time, leaving researchers to focus on data collection and analysis instead of manually counting animals. Notably, citizen science has a considerable impact on participants by allowing thousands of individuals across the globe to learn about animal natural history without having to leave their homes. More specifically, engaging the public in scientific endeavors can help teach scientific literacy and motivate people to make environmentally conscious decisions [37, 38]. Adding a citizen science counting component to drone efforts can help streamline the data collection and interpretation process if the proper precautions are put in place.

One of our critical challenges was determining how to translate multiple citizen scientist counts of each photograph into a single, accurate count. We found that the Median algorithm consistently outperformed the Mean, likely because some extreme counts would have a smaller impact on the Median than they would on the Mean (i.e., the Median value does not depend on the magnitude of all values in the dataset). Unfortunately, algorithms that collapse multiple counts into one cannot provide information on each animal’s location (i.e., x-y coordinates on each photograph) useful for addressing niche partitioning or habitat utilization questions. Such questions can be addressed by comparing expert counts, which are relatively easy but take approximately one workday per drone flight (Wood, *Pers. Obs.*) to six weeks for citizen scientists.

Machine learning algorithms are promising tools for automated counting of pinnipeds in photographs [39, 40]; however, this method is not without challenges [41]. Using this dataset, our team attempted to work with a machine learning company. While total pinniped counts were accurate, the lack of ability to differentiate between species (e.g., seals and sea lions) and age classes (e.g., pups and adults) constrained the questions that could be asked and showed the promise of the citizen science approach.

The accuracy we report may be higher than observed in other mammal or bird species because pinnipeds’ large size and distinctive shapes make them excellent subjects [42]. Additionally, a more expensive drone with a higher resolution camera could be used to increase accuracy; however, these tend to be larger drones with more potential for animal disturbance [43]. Future research should determine whether drones with thermal imaging payloads are better for identifying camouflaged animals or differentiating between grouped animals [44].

## 5. Conclusions

Drones continue to improve as technology becomes more sophisticated [30]. In some cases, drones could replace ground counts entirely and increase the overall accuracy of census data. When combined with strategic citizen science programs, drone imagery can be used to produce accurate data quickly and reduce labor for researchers. We did not quantify this project’s benefit to citizen scientists themselves, but other studies have demonstrated increased ownership and positive environmental attitudes associated with participation [37, 38]. As citizen science becomes increasingly common, it is essential to continue validating each unique project’s data accuracy. Our data suggest that simple tasks with visually interesting photographs, short tutorials, and frequent volunteer interaction opportunities are ideal for engaging citizen scientists. Our project demonstrates that a large-scale, laborious project such as counting pinnipeds can be made impactful and accurate by engaging the public through citizen science. The data that we gathered can be further analyzed for geospatial patterns, range expansion, habitat partitioning, and population changes. We hope that our validation will set the stage for future in-depth analyses of marine mammals that live and breed on islands globally.

## Acknowledgments

We are indebted to the thousands of volunteers that counted animals, including super volunteers Charlotte Lenox, Kristen Cotiaux, Julie Wood, Emily Sullivan, Cormack Pegau, and Helen Bennett. Thank you to Drs. Erika Zavaleta and Judy Straub for making this project possible; to the staff at Zooniverse, especially Cliff Johnson and Grant Miller, for their guidance in project design; to Katie Sweeney and Jen Cormier for their expertise with website design and volunteer recruitment; to Abram Fleishman for creating and sharing image manipulation code; and to past and present members of the Costa lab and Año Nuevo State Park staff and docents. Our research was completed at the University of California Natural Reserve System’s Año Nuevo Reserve. The research was approved by the University of California Santa Cruz Institutional Animal Care and Use Committee #Costd1709 and the National Marine Fisheries Service marine mammal research permit #19108. Drone flights were authorized by the University of California Center of Excellence for Unmanned Aircraft Systems Safety with permission from Año Nuevo State Park. This publication uses data generated via the Zooniverse.org platform funded by generous support from a Google Global Impact Award and from the Alfred P. Sloan Foundation. Financial support for this research was provided by the Friends of the Seymour Marine Discovery Center Student Research and Education Award (to SAW), the Center to Advance Mentored, Inquiry-Based Opportunities in Ecology and Conservation (to SAW), the Packard Ocean Science and Technology Endowment (to RSB and DPC), and a National Science Foundation Postdoctoral Research Fellowship in Biology (to RSB).

